# Errors-in-variables and validation problems in reaction norm predictions for wild populations

**DOI:** 10.1101/2024.12.30.630729

**Authors:** Rolf Ergon

## Abstract

Studies of phenotypic responses in wild populations are often based on reaction norm models where the environmental drivers in many cases are related to climate change. Such input signals will never be exactly known, and there will always be measurement errors also in the recorded responses. In parameter estimation these errors give rise to errors-in-variables problems, especially in form of overfitting caused by errors in the input measurements. A second important feature of such phenotypic response problems is that the environmental inputs must be given appropriate but largely unknown reference values. A third problem is that it is difficult to find good validation methods for predicted responses. Essential aspects of these problems are here studied by use of a reaction norm model in its simplest univariate form, characterized by a mean intercept value and a mean plasticity slope value, and the overall conclusion is that validated disentanglement of plasticity and genetic adaptation based on realistically short data for wild populations is a difficult task. In a proposed validation method, the available input-output data is split into one part for modeling and one part for validation, and the feasibility of this approach is studied in simulations with use of a prediction error method, which is essentially a maximum likelihood method. It is also argued that validation of a chosen or estimated reference environment in practice is impossible when the data comes from the (unintended) anthropogenic global warming experiment, where no independent experimental data exists. When the evolution is slow because of small genetic variances, overlapping generations and long lifetimes, or because of near optimal adaptive plasticity, the best quantitative genetics option may be to assume a constant plasticity slope value, equal to the initial value. It turns out to be easy to estimate this value, but that should be done without setting other unknown parameter values to zero. This option is appealing also because it removes the dependence of a guessed or estimated reference environment.

## 1 Introduction

Errors-in-variables problems are ubiquitous in ecological and evolutionary modeling where processes driven by environmental variables are modeled for some specific purpose. For example, Kangas (1998) used tree diameter at breast height, tree height and the height to the base of live crown as independent variables in a forest growth model and discussed the effects of measurement errors. Stoklosa et al. (2015) discussed how measurement errors in spatial climate variables affect species distribution models. A number of articles have studied effects of climate change and discussed whether phenotypic changes are genetically based or the result of phenotypic plasticity. A summary of such studies was given in Merilä and Hendry (2014), with reference to 11 accompanying studies covering a variety of taxa. A main conclusion was that plastic responses on climate change are common, but that there were few examples of adaptive genetic responses. An early example where adaptive responses were found was a study of a northern squirrel population (Réale et al., 2003), but that was for the special case with a constant plasticity slope. In another example Tarka et al. (2015) studied great reed warblers, while Bonamour et al. (2019) used plasticity of blue tit breeding phenology as an example. A common feature in such papers is the use of various ecological drivers of evolutionary processes, and such drivers are obviously not exactly known. In a recent example, Valdés et al. (2023) used 22 years of field observations of the perennial forest herb *Lathyrus vernus* in order to assess phenotypic selection of flowering time as function of spring temperature, and as spring temperature they used the mean April temperature as an approximation of the true but unknown driver.

In all these examples, and numerous others, there are errors-in-variables problems as discussed in Söderström (2019) and depicted in Fig. 1. The ecological system has scalar or vector valued and time-varying input and output signals *u*_0,*t*_ and *z*_0,*t*_, respectively, while the measured signals are *u*_*t*_ = *u*_0,*t*_ + *ũ*_*t*_ and 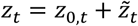, respectively, where *ũ*_*t*_ and 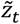 are the measurement errors. Without further assumptions there is a fundamental lack of identifiability in the errors-in-variables problem.

**Figure 1.**
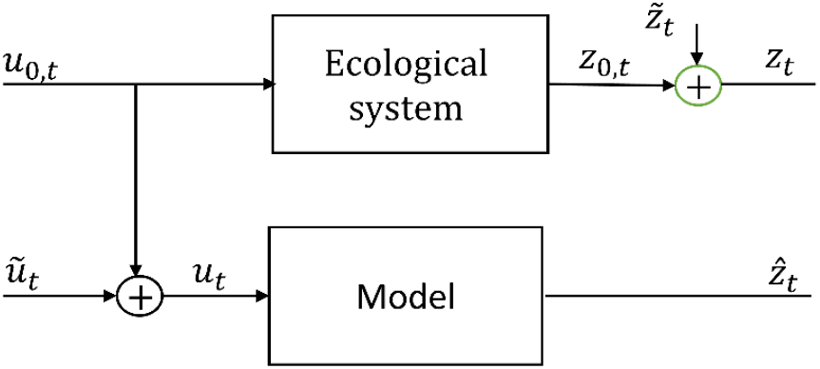
The basic setup for an errors-in-variables model identification problem.

In the ecological and evolutionary setting the true output *z*_0,*t*_ is typically multivariate, with vectors ***y***_0,*t*_ and ***w***_0,*t*_ of individual phenotypic and relative fitness values, respectively. The true input *u*_0,*t*_ that drives the evolutionary process is often a climatic variable, which varies from year to year with very little autocorrelation. In such cases the influence of the environmental cue *u*_0,*t*_ is found from *u*′_0,*t*_ = *u*_0,*t*_ − *u*_*ref*_, from which follows *u*′_*t*_ = *u*_*t*_ − *u*_*ref*_, where *u*_*ref*_ is a reference environment as discussed in Ergon (2022). An important climatic variable is spring air temperature, which typically is a Gaussian white noise process around a slow trend in mean value, as shown in Fig. 2. Although the specific choices of such input signals as drivers of evolutionary processes should be based on all available information, they will obviously be more or less error corrupted representations *u*_*t*_ of the true driver *u*_0,*t*_. If also the error signal *ũ*_*t*_ is Gaussian white noise, there is no practical possibility to reconstruct *u*_0,*t*_ from known values of *u*_*t*_. A theoretical possibility could otherwise involve the use of information on non-normal distributions of *u*_0,*t*_ and *ũ*_*t*_ (Söderström, 2019). Without such additional information one must simply accept parameter estimation errors caused by *ũ*_*t*_, and as far as possible test the results by validation against data that are not used in the modeling. A primary validation problem is here that data typically come from field observations of the (unintended) anthropogenic global warming experiment, and that no independent experimental data exist. A possible approach for this type of data is to a use a part of the recorded time series for modeling and the rest for validation. There is also a second validation problem with such data in that the reference environment is unknown, and this problem cannot be solved by splitting the time series data in two parts (because the two parts will have the same reference environment). The unknown reference environment thus represents a serious problem.

**Figure 2.**
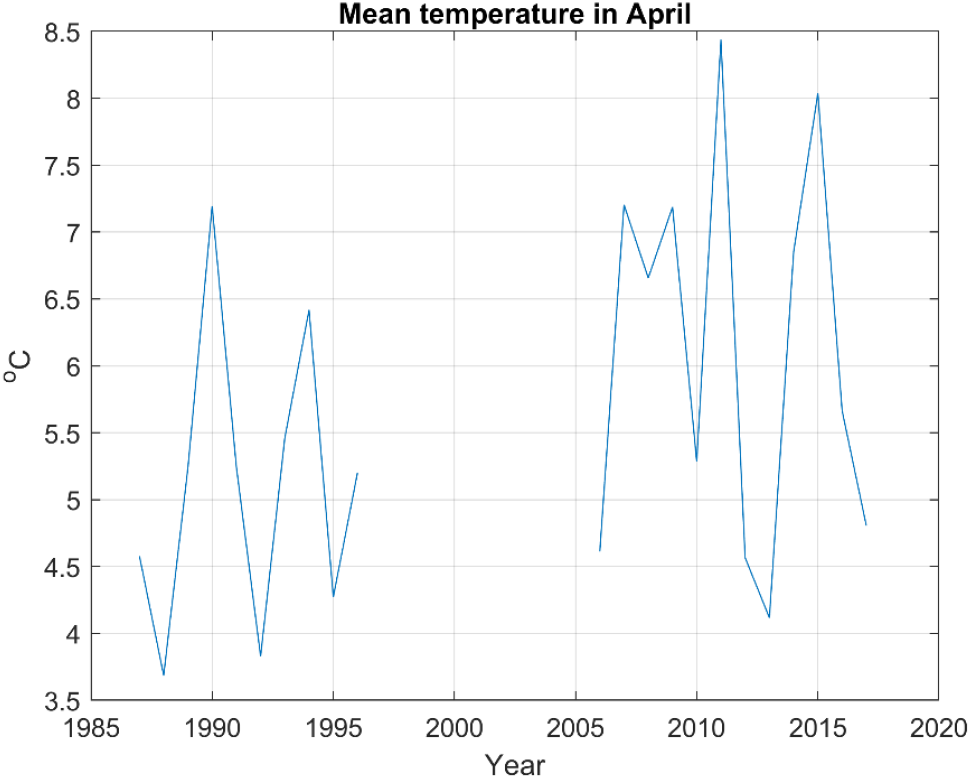
Mean April temperature in Tullgarn south of Stockholm, which is used as environmental driver in Valdés et al. (2023).

Here, I will focus on the common cases where reaction norms are predicted on the basis of time-varying environmental variables, as discussed above and exemplified in Fig. 2. A reaction norm describes the phenotypes that a given set of genes in an individual biological organism can produce across a range of environments. Mean reaction norms in a population of such organisms can evolve by natural selection for example after a sudden introduction to a new environment or as response on climate change, and this evolution can be modeled by use of several methods. In its simplest univariate form, a mean reaction norm is characterized by a mean intercept value and a mean plasticity slope value, but it may also be necessary to use models with a multiple of reaction norms. In many cases reaction norms are also nonlinear (Arnold et al, 2019).

A simple reaction norm model is the two-trait intercept-slope model as used in Lande (2009),

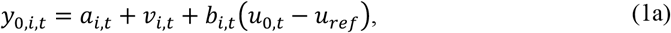

where *y*_0,*i*,*t*_ is the individual phenotypic response on an environmental cue *u*_0,*t*_− *u*_*ref*_ at generation *t*, and where *u*_*ref*_ is the reference environment (which Lande assumed to be zero). In populations with sexual reproduction, an individual will here be a midparent with traits that are mean values of the traits of the two parents. The reference environment is defined as the environment where the phenotypic variance has its minimum, and where thus the expected geometric mean fitness is maximized (Ergon, 2022). In the following I will treat the individual reaction norm parameters *a*_*i*,*t*_+ *v*_*i*,*t*_ and *b*_*i*,*t*_ as quantitative traits in their own right, and where necessary I will thus distinguish between phenotypic traits and reaction norm traits. In eq. (1a), *a*_*i*,*t*_ and *b*_*i*,*t*_ are the additive genetic components of the reaction norm traits, while *v*_*i*,*t*_ is zero mean Gaussian white noise. From eq. (1a) follows the mean reaction norm equation

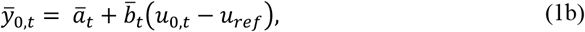

where the mean traits *ā*_*t*_ and 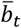, and thus also 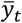, may evolve as results of a changing environment and natural selection. See Lande (2009) for step response simulations of the model in eqs. (1a,b), and Fig. 1 in Ergon (2023b) for a detailed explanation.

As discussed in Ergon (2023b), it may be difficult to estimate the environmental reference values from data, and I will therefore assume that *u*_*ref*_ is known from historical data, for example from the time before the recent increase in global temperature, or before a sudden change in the environment. In simulations, I will assume that *u*_*t*_ is a temperature, and I will then choose a temperature scale such that *u*_*ref*_ = 0.

A main focus in the presentation will be to show the difficulties involved in disentanglement of the plasticity and microevolutionary contributions to measured changes in mean phenotypic values, given short time series data collected in field studies, where also the reference environment is unknown. For this purpose I will use simulations where assumed true data are generated according to the model in eq. (1a,b), and where parameters in this model are found by use of a prediction error method (PEM). The PEM approach is well established in the engineering control field (Ljung, 1998; Ljung, 2010), but although comparisons between measured and predicted responses are discussed in the literature (Morrissey et al., 2010; Merilä et al., 2014), PEM as such appears to be largely unknown in the quantitative genetics community. A main parameter estimation difficulty is that errors in the assumed environmental drivers will result in large prediction errors caused by overfitting, and this difficulty becomes even more pronounced when a part of the already short data is set aside for validation purposes. The situation may be different in controlled laboratory settings, where measurement errors may be small, and where the time series may be longer. It should be noted that overfitting to short data is not a specific PEM problem, errors in the assumed environmental driver will result in overfitting regardless of the choice of parameter estimation method.

Background theory is given in Section 2, together with a more detailed discussion of the input measurement problem and a presentation of the proposed validation methodology under the assumption of a correct reference environment. A special case with constant plasticity slope is described. Different aspects of the identification and validation processes are illustrated and documented by simulations in Section 3. In the simulations I assume the simple model (1a,b), with parameter values as in Lande (2009), and I use two types of environmental changes. The first type is a sudden change in environment, as in Lande (2009), while the other type is a gradual change similar to the global warming since 1970. In both cases I assume non-overlapping generations and realistic temperature variations from generation to generation, but extensions to cases with overlapping generations are also discussed. A final discussion with conclusions is given in Section 4. Additional examples of prediction results are given in Appendices A and B. Typical MATLAB codes are given in Appendix C.

## 2 Theory and Methods

### 2.1 The selection gradient model

Under the assumption of random mating in a large population the evolution of correlated phenotypic traits is determined by the multivariate breeder’s equation (Lande, 1979). When the reaction norm parameters in eq. (1a) are treated as quantitative traits in their own right, the multivariate breeder’s equation gives

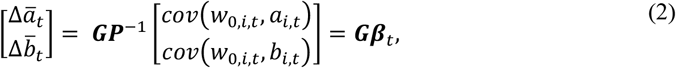

where ***β***_*t*_ is the selection gradient. Here, 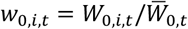 is the relative individual fitness, where *W*_0,*i*,*t*_ and 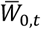 are the individual and mean fitness, respectively, while ***G*** and ***P*** are given by 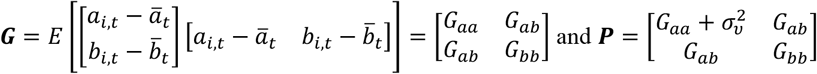. I will assume that the ***G*** and ***P*** matrices are constant.

The model in eq. (2) cannot be identified by use of available environmental, phenotypic, and fitness data, for the simple reason that the individual latent variables *a*_*i*,*t*_ and *b*_*i*,*t*_ are not available. This problem was solved in Ergon (2022) by use of a linear transformation of the vector [*a*_*i*,*t*_ *b*_*i*,*t*_]^*T*^ onto the vector [*a*_*i*,*t*_ *b*_*i*,*t*_ *y*_*i*,*t*_]^*T*^. This results in the model

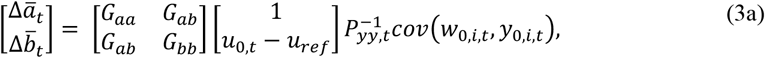

where

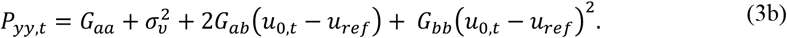

In eq. (3a), selection with respect to *a*_*i*,*t*_ and *b*_*i*,*t*_, as in eq. (2), is replaced by selection with respect to *y*_0,*i*,*t*_, which is assumed to be known (with a measurement error). The two models (2) and (3a,b) give equal results when the population size *N* → ∞, but for practical purposes they are interchangeable also for rather limited population sizes. Since both models in any case are approximations of reality, eqs. (3a,b) can be seen as just as correct as eq. (2). Note that eq. (3a) is a dynamical model in the sense that *P*_*yy*,*t*_ varies with time. In order for eq. (3a) to be compatible with the fundamental Price equation (Price, 1970; Ergon, 2019) the covariance expression should be computed as

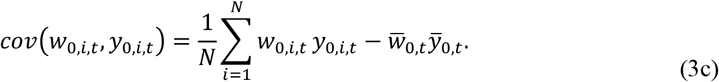

With given initial mean trait values *ā*_1_ and 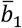, eq. (3a) gives *ā*_*t*_ and 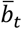, and thus also 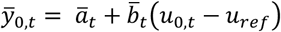 according to eq. (1b), for use as assumed ‘true’ mean phenotypic data in simulations.

### 2.1 The prediction error model

Under the assumption of a true model according to eqs. (3a,b), and with measurement errors, the prediction error method (PEM) uses the model

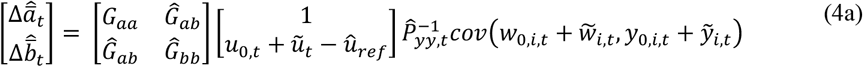

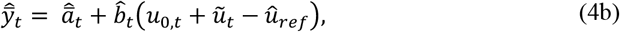

where

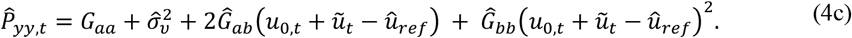

In addition, PEM uses the unknown initial mean reaction norm slope value 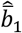, while the initial intercept value is given by 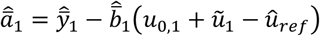 Here, 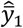 can be set to any value, for example 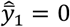. Note that with 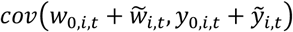 given from the data, *G*_*aa*_ can be set to any value, such that and *Ĝ*_*bb*_ and 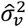 are found in relation to this value, i.e., *G*_*aa*_ cannot be estimated.

Eq. (3) is derived under the assumption of non-overlapping generations. With overlapping generations only a fraction *c*_*t*_ < 1 of the population is affected by the changes in fitness, and the expression 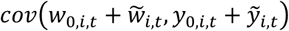 in eq. (4a) should then be replaced by *c*_*t*_ 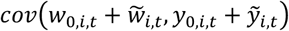 (Ergon, 2023a). A constant value of *c*_*t*_ can also be estimated from input-output data.

In PEM, all the unknown parameters *Ĝ*_*ab*_, *Ĝ*_*bb*_, 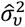, *b̂*_1_ and *û*_*ref*_ (and possibly *c*_*t*_) are in principle tuned until the criterion function 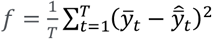 reaches a minimum, where *T* is the number of samples. In the simulations in Section 3 the MATLAB function *fmincon* is used for this purpose. This function finds a constrained minimum of a scalar function of several variables. In order to focus on the errors-in-variables and validation problems, the simulations assume that *Ĝ*_*ab*_ = *G*_*ab*_ = 0 and *u*_*ref*_ = 0. As will be found in the simulations, the final residuals 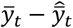 will approximately be white noise, which makes PEM into a maximum likelihood method (Ljung, 2010).

### 2.2 PEM error analyses

Since model tuning results in 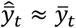, all measurement errors must be compensated by errors in the estimated parameters, including 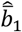 and possibly *û*_*ref*_ and *c*_*t*_. As an analysis of how combinations of measurement errors affect the results becomes quite unwieldy, I here discuss the effects of such errors one at a time.

#### 2.2.1 Errors in *û*_*ref*_

Theory and simulations in Ergon (2022) show that errors in *û*_*ref*_ give large errors in the predicted mean reaction norms, and with perfect measurements and known parameter values it is theoretically possible to compute these errors. In practice, errors in *û*_*ref*_ may give large errors in the predicted mean reaction norms, as shown in simulations in Section 3.

#### 2.2.2 Measurement errors in *y*_*i*,*t*_ and/or *w*_*i*,*t*_, with *u*_*ref*_ known

For large populations, white noise errors 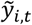 and 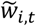 will neither affect 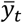 nor *cov*(*w*_*i*,*t*_, *y*_*i*,*t*_), and will thus have no effects on the mean trait predictions. For small population sizes, however, such random errors will not be zero mean, and 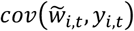 may not be quite zero, and simulations will show some effects on the predictions.

#### 2.2.3 Measurement errors in *u*_*t*_, with *u*_*ref*_ known

The main problem with measurement errors *ũ*_*t*_ as shown in Fig.1 is that overfitting may give large errors in the estimated parameter values. The true input *u*_0,*t*_ is typically a white noise process added to a slow trend, as shown in Fig. 2. With white noise measurement errors *ũ*_*t*_ it will then in practice be impossible to separate the effects of *u*_0,*t*_ and *ũ*_*t*_ from each other, and there is therefore no option but to live with the overfitting problems caused by *ũ*_*t*_. These problems are especially pronounced with short data. The underlying problem is that minimization of 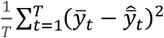 does not imply minimization of 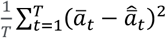 and 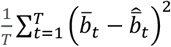, unless both the model and the measurements are perfect. This will be demonstrated in simulations in Section 3.

### 2.3 The model validation problem

In addition to errors-in-variables problems, a second problem is validation of the model found by use of the PEM approach, and thus of the predicted mean reaction norm traits over time. This validation problem is not specific for PEM, i.e., there will be an errors-in-variables problem independent of parameter estimation method.

The typical industrial approach is to test the model against a new and independent set of input-output data (Ljung, 2010), and this is also essential for predictive models in ecology (Tredennic et al., 2021). This is obviously not possible for a model of reaction norm responses on climate change, where time series of the environmental variable *u*_*t*_ are given by a single natural experiment. Under the assumption of a correct reference environment, model validation can still be performed by splitting the time series into two parts, one for modeling and one for validation, as further discussed below. However, validation of a guessed or estimated reference environment, i.e., of which environment the population is adapted to, will require comparison with data from a population that has evolved from an adapted state in a different environment, and this is not a feasible approach for wild populations. The proposed validation method therefore assumes that the guessed reference environment is correct. The validation process has three steps, using the model in eqs. (1a,b) as example. First, identify an initial model by use of for example the first 3/4 of the generations, and find predictions 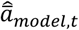 and 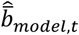 for all generations, including the remaining 1/4 of the generations. Second, identify a final model by use of all data, and find predictions 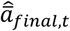 and 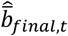 for all generations. Third, compare the predicted mean trait changes over all generations by use of the initial model, 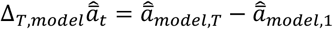 and 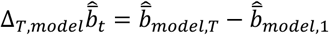, with the same changes computed by use of the final model, i.e., with 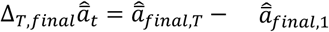 and 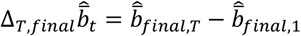. For a good model this comparison will result in validation ratios 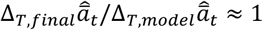 and 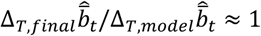, but a possible conclusion could be that the predictions 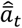 can be trusted, while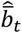 cannot. This possibility can be seen by a theoretical study of the changes 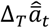 and 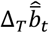 over all samples as functions of Δ_*T*_ *u*_*t*_ = *u*_*T*_ − *u*_1_. From eq. (1b) follows

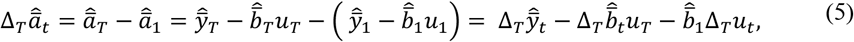

which shows that we with 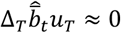, and a good estimate 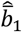, may find a good prediction 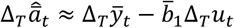 also if the prediction 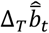 is poor.

### 2.4 Special case with *G*_*bb*_ = 0

For the special case with *G*_*bb*_ = 0, eq. (4) as well as eq. (1b) gives

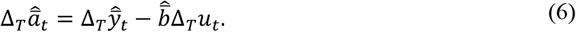

We then have what might be called constant partially adaptive plasticity (Lande, 2009), i.e., the constant plasticity plays a role in the evolution of *ā*_*t*_ and 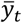 It follows from eq. (6) that it in this case is not necessary to know *u*_*ref*_ in order to find the predicted change in *ā*_*t*_ and 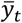 over all generations.

## 3 Simulations

### 3.1 Description of simulation system

As in Lande (2009), assume the reaction norm model (1a,b) with *u*_*ref*_ = 0, i.e., with true mean phenotypic and reaction norm traits 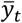, *ā*_*t*_ and 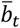. True responses are generated by the model (3a,b), with *G*_*aa*_ = 0.5, *G*_*bb*_ = 0.045, *G*_*ab*_ = 0, and 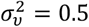. Also assume non-overlapping generations where the individual values *a*_*i*,*t*_, *b*_*i*,*t*_ and *v*_*i*,*t*_ at each generation are drawn from populations with normal distributions around *ā*_*t*_,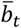 and 0, respectively. The individual fitness function is assumed to be

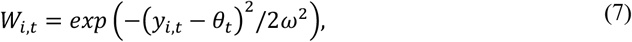

where *θ*_*t*_ is the stochastic phenotypic value that maximizes fitness, while *ω*^2^ = 50. We may for example assume that *θ*_*t*_ is the optimal breeding time as function of spring temperature.

Also assume a stochastic environment *u*_*t*_, with mean *μ*_*U*,*t*_, and added zero mean Gaussian and white variations *u*_*n,t*_ with variance 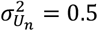, i.e., *u*_*t*_ = *μ*_*U*,*t*_+ *u*_*n,t*_. The population is assumed to be fully adapted to a stationary stochastic environment with mean value *μ*_*U*,*t*_ = *u*_*ref*_ = 0. In a corresponding way assume that *θ*_*t*_ = *μ*_Θ,t_+ *θ*_*n,t*_, where *θ*_*n,t*_ is zero mean Gaussian and white with variance 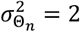, and where *u*_*n,t*_ and *θ*_*n,t*_ are correlated with covariance 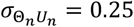. Following Lande (2009), we may assume that juveniles of generation *t*are exposed to the environment *u*_*t*−*τ*_ during a critical period of development a fraction of a generation before the adult phenotype is expressed and subjected to natural selection. Define *θ*_*t*_ = 2*u*_*t*_, which implies a linear relationship 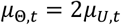, variances 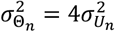, and covariance 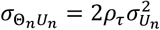, where *ρ*_*τ*_ = 0.25 is the autocorrelation of background environmental fluctuations. The optimal value of the mean plasticity slope in a stationary stochastic environment is then 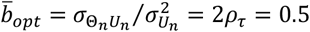 (Lande, 2009), and this value is in the simulations used as initial mean slope value, i.e., 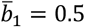. Since 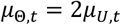, this means that the population in a stationary stochastic environment has 25% adaptive plasticity.

### 3.2 Two test cases, with step and ramp inputs

Simulation results are given for two different cases, with input signals as shown in Fig. 3. In the first case the mean environment *μ*_*U*,*t*_ is a step function from 0 to 2.5 at *t*= 10, while it in the second case is a ramp function where *μ*_*U*,*t*_ goes from 0 to 2.5 over 50 generations. The noisy ramp function is similar to the registered yearly mean temperatures in Oslo, Norway, from 1970 to 2020 (Norsk klimaservicesenter), using the temperature around 1960 as zero-point.

**Figure 3.**
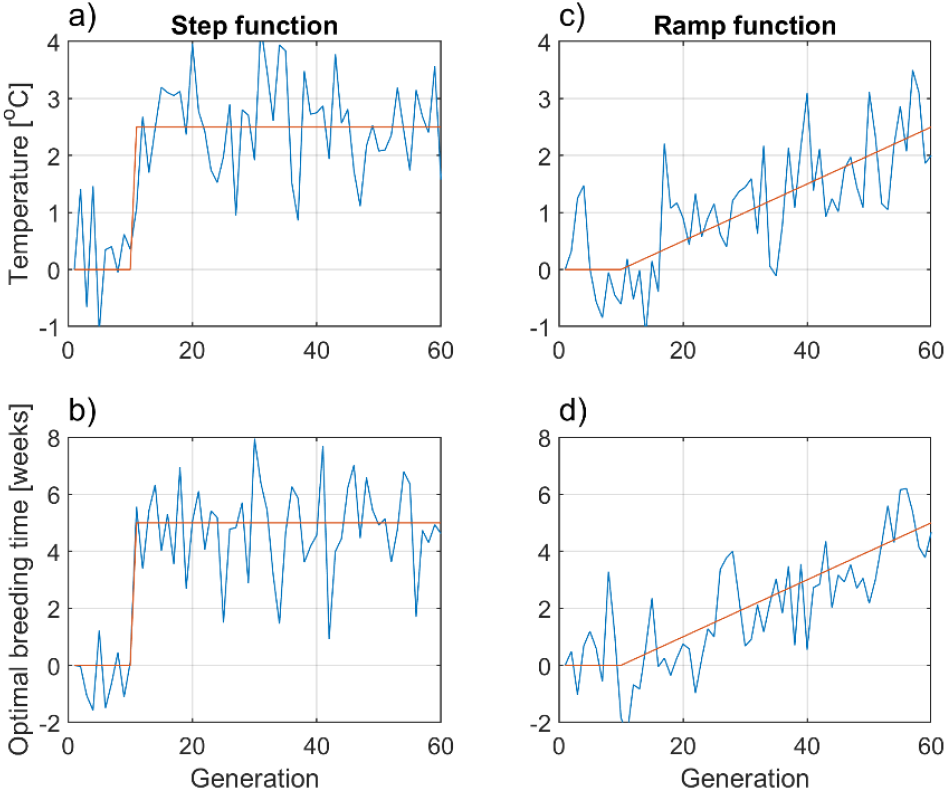
Input signals in the form of noisy step and ramp functions, with variances 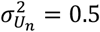 and 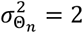, and covariance 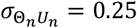. Panels b) and d) show advancements in breeding time *θ*_*t*_ towards earlier dates as results of increased temperatures.

### 3.3 Effects of random input measurement errors

As discussed in Subsection 2.2, random errors in the measured input signal *u*_*t*_ implies an error-in-variables problem, resulting in errors in the predicted mean reaction norm traits. The problem is that the estimated parameters *Ĝ*_*bb*_ and 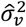 are obtained by overfitting of 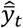, which results in errors in the predicted mean traits 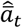 and 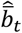, as shown in Fig. 4. Here, recall that we in eq. (4a) can set *G*_*aa*_ to any fixed value and scale *Ĝ*_*bb*_ and 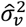 accordingly. The plots in Fig. 4 also show that the very same errors in the input signal that cause large errors in especially *Ĝ*_*bb*_, and thus in the predicted reaction norm traits, have less influence on the predictions when a model with true parameter values is used. For clarity it is assumed that the initial mean plasticity slope value 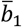 is known, such that PEM only has to find *Ĝ*_*bb*_ and 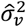. The prediction errors will be larger when also 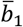 must be estimated.

**Figure 4.**
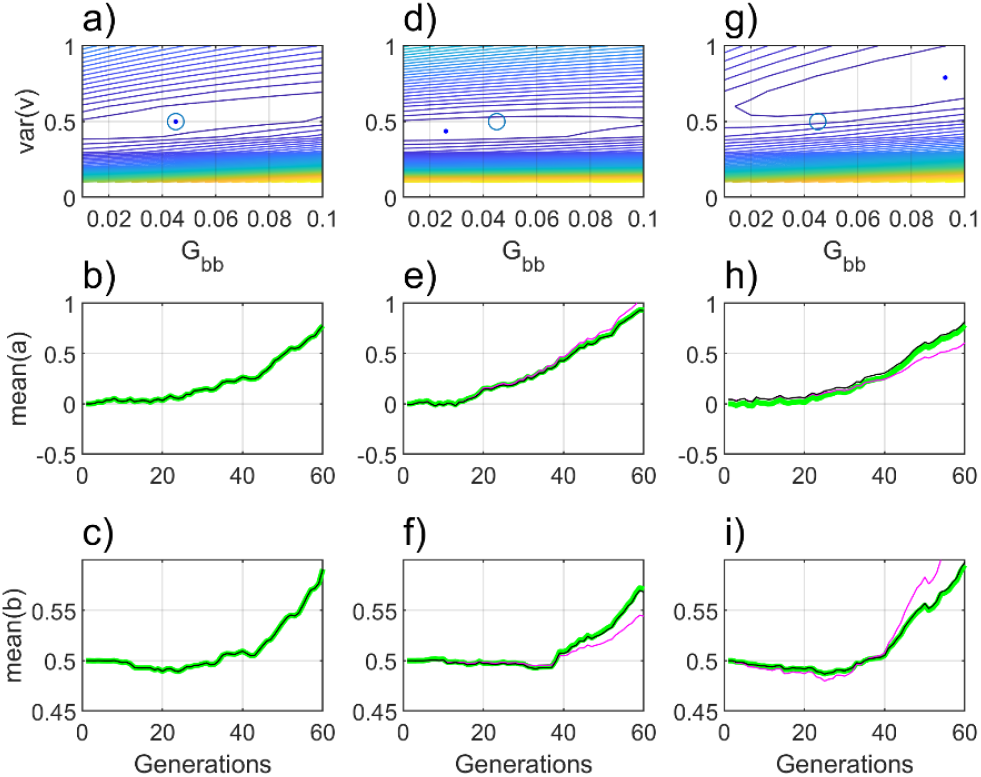
Contour plots with true (circles) and estimated (dots) parameter values (upper panels). Here, the x and y variables are *G*_*bb*_ and 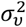, while the dependent variable is 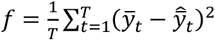 as function of *G*_*bb*_ and 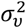. The corresponding true and predicted mean reaction norm traits are shown in the lower panels, as green and red lines, respectively. With perfect input and output measurements, the estimated parameter values and predicted responses are also perfect as shown in panels a), b) and c). Random errors in the measured input *u*_*t*_ result in errors in the estimated parameter values and the predicted mean traits, as shown for two realizations in panels d) to i). The white measurement errors had standard deviations that were 20% of the standard deviation of the random variations in *u*_*t*_. For the two tuning results marked by dots in panels d) and g), the minimum values of 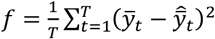 were 25 · 10^−5^ and 35 · 10^−5^, respectively. Response plots with the very same inputs applied on a model with true parameter values (circles in the contour plots) are shown by black lines, corresponding to *f* = 29 · 10^−5^ and *f* = 52 · 10^−5^, respectively, i.e., larger than the optimal minimum values, but still resulting in clearly better predictions.

### 3.4 PEM identification and validation with correct model structure

#### 3.4.1 Results with *u*_*ref*_ known

The true system was simulated with a population of *N* = 1,000 individuals over *T* = 60 generations (increased to *T* = 120 in two cases). Individual measurement errors 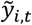 and 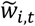 were introduced as Gaussian white noise with standard deviations that were 20% of the registered mean standard deviations of *y*_0,*i*,*t*_ and *w*_0,*i*,*t*_ over all generations, respectively. Measurement errors *ũ*_*t*_ were introduced as Gaussian white noise with standard deviation that was 20% of the standard deviation of the known random variations in *u*_*t*_. Identification and validation were performed in three steps, as described in Subsection 2.3. First, data for the first 3T/4 generations (modeling set) were used for identification of a preliminary model, which was used to find predictions 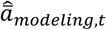 and 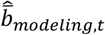 for all generations, including the last *T*/4 generations. Second, a final model was found by use of all data, which gave predictions 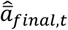 and 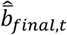 . Third, validation ratios 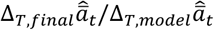 and 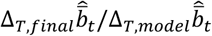 were computed, where Δ_*T*_ stands for change over *T* = 60 generations. Finally, the predicted changes in mean traits were compared with the true changes, i.e., 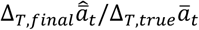 and 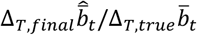 were computed. The lower and upper search limits for 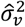 were set to 0.5*G*_*aa*_ and 2*G*_*aa*_, respectively, which means that the limits for the heritability 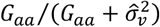 were 0.33 and 0.67.

Results are given in Tables 1 and 2, presented as *mean* ± *SE* values from 100 repeated simulations with different realizations of all random variables. The starting values in the numerical searches were set to 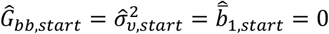. As pointed out in Subsection 2.3, good and validated predictions are indicated by 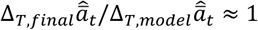 and 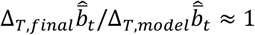. Perfect measurements of *u* results in good predictions (Cases 1 and 4), while measurement errors in *u*_0,*t*_ combined with short data gives poor predictions of especially 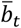(Cases 2 and 5). Note that the changes in the mean trait *ā*_*t*_ in Cases 2 and 5 were clearly overestimated. Also note that the predictions were improved when *T* was increased from *T* = 60 to *T* = 120 (Cases 3 and 6), especially in the ramp responses, but that such an increase is unrealistic for many wild populations.

**Table 1.** Estimation and prediction results for system given by eqs. (3a,b) and (7), and use of the step function in Fig. 3, panels a) and b). The reference environment *u*_*ref*_ = 0 is assumed to be known. Case 1: Step function, with measurement errors in *y*_0,*i*,*t*_ and *w*_0,*i*,*t*_. Case 2: Step function, with measurement errors in *y*_0,*i*,*t*_, *w*_0,*i*,*t*_ and *u*_0,*t*_. Case 3: Same as Case 2, but with *T* = 120.

| Parameters and prediction ratios | True or optimal values | Case 1 | Case 2 | Case 3 |
| --- | --- | --- | --- | --- |
| $\hat{G}_{bb}$ | 0.045 | $0.045 \pm 0.001$ | $0.036 \pm 0.011$ | $0.040 \pm 0.011$ |
| $\hat{\sigma}_v^2$ | 0.5 | $0.50 \pm 0.01$ | $0.45 \pm 0.06$ | $0.47 \pm 0.06$ |
| $\hat{b}_1$ | 0.5 | $0.50 \pm 0.00$ | $0.50 \pm 0.02$ | $0.49 \pm 0.04$ |
| $f_{final}$ | 0 | $(51 \pm 13)10^{-6}$ | $(83 \pm 18)10^{-4}$ | $(40 \pm 6)10^{-4}$ |
| $\Delta_{T,true} \bar{a}_t / \Delta_{T,true} \bar{y}_t$ | - | $0.40 \pm 0.08$ | $0.41 \pm 0.08$ | $0.54 \pm 0.23$ |
| $\Delta_{T,true} \bar{b}_t / \bar{b}_1$ | - | $0.56 \pm 0.04$ | $0.57 \pm 0.04$ | $0.68 \pm 0.04$ |
| $\Delta_{T,final} \hat{\bar{a}}_t / \Delta_{T,model} \hat{\bar{a}}_t$ | 1 | $1.00 \pm 0.01$ | $0.98 \pm 0.08$ | $1.00 \pm 0.03$ |
| $\Delta_{T,final} \hat{\bar{b}}_t / \Delta_{T,model} \hat{\bar{b}}_t$ | 1 | $1.00 \pm 0.02$ | $1.10 \pm 0.24$ | $1.00 \pm 0.09$ |
| $\Delta_{T,final} \hat{\bar{a}}_t / \Delta_{T,true} \bar{a}_t$ | 1 | $1.00 \pm 0.01$ | $1.10 \pm 0.11$ | $1.05 \pm 0.10$ |
| $\Delta_{T,final} \hat{\bar{b}}_t / \Delta_{T,true} \bar{b}_t$ | 1 | $0.99 \pm 0.02$ | $0.85 \pm 0.17$ | $0.92 \pm 0.16$ |

**Table 2.** Estimation and prediction results for system given by eqs. (3a,b) and (7), and use of the ramp function in Fig. 3, panels c) and d). The reference environment *u*_*ref*_ = 0 is assumed to be known. Case 4: Ramp function, with measurement errors in *y*_0,*i*,*t*_ and *w*_0,*i*,*t*_. Case 5: Ramp function, with measurement errors in *y*_0,*i*,*t*_, *w*_0,*i*,*t*_ and *u*_0,*t*_. Case 6: Same as Case 5, but with *T* = 120 and *μ*_*U*,*t*_ = 2.5 for *t* > 60.

| Parameters and prediction ratios | True or optimal values | Case 4 | Case 5 | Case 6 |
| --- | --- | --- | --- | --- |
| $\hat{G}_{bb}$ | 0.045 | $0.045 \pm 0.004$ | $0.039 \pm 0.030$ | $0.041 \pm 0.009$ |
| $\hat{\sigma}_v^2$ | 0.5 | $0.50 \pm 0.02$ | $0.42 \pm 0.11$ | $0.44 \pm 0.07$ |
| $\hat{b}_1$ | 0.5 | $0.50 \pm 0.00$ | $0.48 \pm 0.02$ | $0.48 \pm 0.03$ |
| $f_{final}$ | 0 | $(52 \pm 12)10^{-6}$ | $(73 \pm 37)10^{-4}$ | $(42 \pm 10)10^{-3}$ |
| $\Delta_{T,true} \bar{a}_t / \Delta_{T,true} \bar{y}_t$ | - | $0.35 \pm 0.09$ | $0.34 \pm 0.21$ | $0.30 \pm 0.06$ |
| $\Delta_{T,true} \bar{b}_t / \bar{b}_1$ | - | $0.21 \pm 0.03$ | $0.21 \pm 0.03$ | $1.21 \pm 0.04$ |
| $\Delta_{T,final} \hat{\bar{a}}_t / \Delta_{T,model} \hat{\bar{a}}_t$ | 1 | $0.99 \pm 0.03$ | $1.06 \pm 0.18$ | $1.01 \pm 0.10$ |
| $\Delta_{T,final} \hat{\bar{b}}_t / \Delta_{T,model} \hat{\bar{b}}_t$ | 1 | $1.03 \pm 0.19$ | $351 \pm 1357 *$ | $1.03 \pm 0.16$ |
| $\Delta_{T,final} \hat{\bar{a}}_t / \Delta_{T,true} \bar{a}_t$ | 1 | $1.00 \pm 0.02$ | $1.13 \pm 0.21$ | $1.09 \pm 0.12$ |
| $\Delta_{T,final} \hat{\bar{b}}_t / \Delta_{T,true} \bar{b}_t$ | 1 | $0.99 \pm 0.07$ | $0.85 \pm 0.52$ | $0.97 \pm 0.10$ |

\*Some values of $\Delta_{T,model} \hat{\bar{b}}_t$ are close to zero.

The system in eqs. (3a,b) and (7), with given parameter values and ramp inputs as in Fig. 3, gave what is considered to be true mean trait responses as shown in Fig. 5, where also typical predicted responses are shown. Fig. 5 also shows to which degree the predicted responses 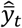 are similar to the true responses 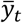, which is a prerequisite for useful predictions of the mean reaction norm traits. Panel d) in Fig. 5 shows for example that a model based on samples 1 to 45 gives a poor fit for the validation samples 46 to 60. The autocorrelation plot in Fig. 6 shows that typical final residuals 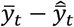 essentially are white noise sequences, which makes PEM into a maximum likelihood method (Ljung, 2010).

**Figure 5.**
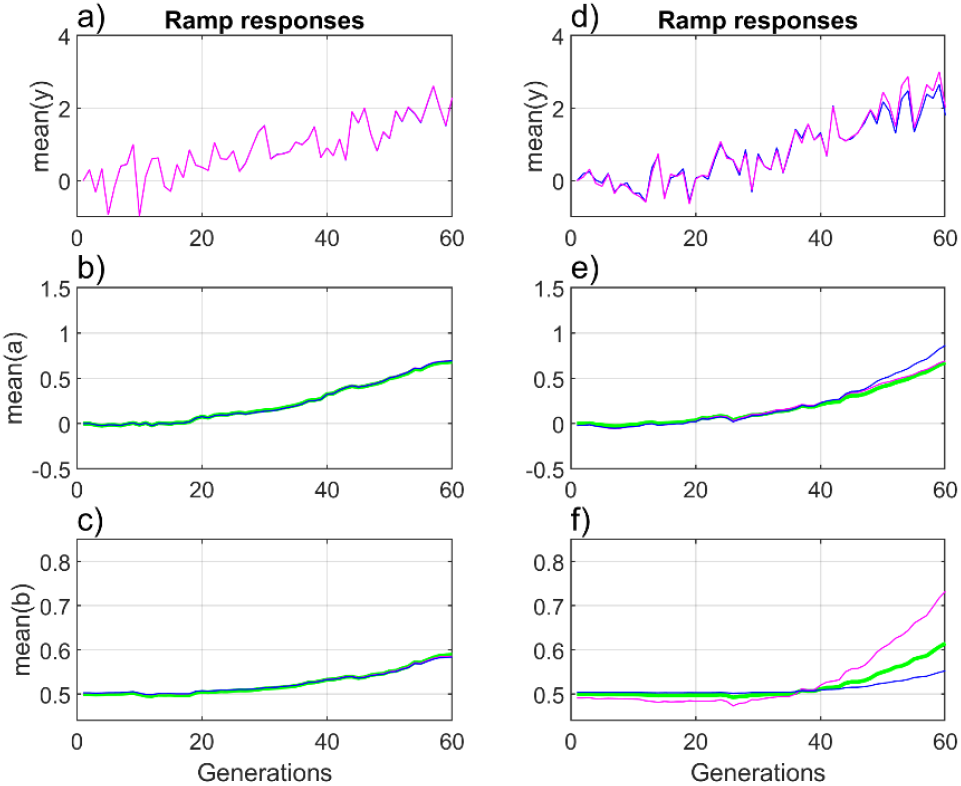
Typical responses on the noisy ramp function as shown in Fig. 3. Responses with measurement errors in *y*_0,*i*,*t*_ and *w*_0,*i*,*t*_ are shown in panels a), b) and c), and responses with measurement errors also in *u*_0,*t*_ are shown in panels d), e) and f). Panels a) and d) show 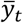 (blue lines) and 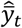 based on the modeling data (red lines). Panels b) and e) show the true responses *ā*_*t*_ (green lines), and the predicted responses 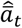 based on the modeling data (red lines) and on all data (blue lines). Panels c) and f) show the true responses 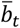 (green lines), and the predicted responses 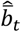 based on the modeling data (red lines) and on all data (blue lines).

**Figure 6.**
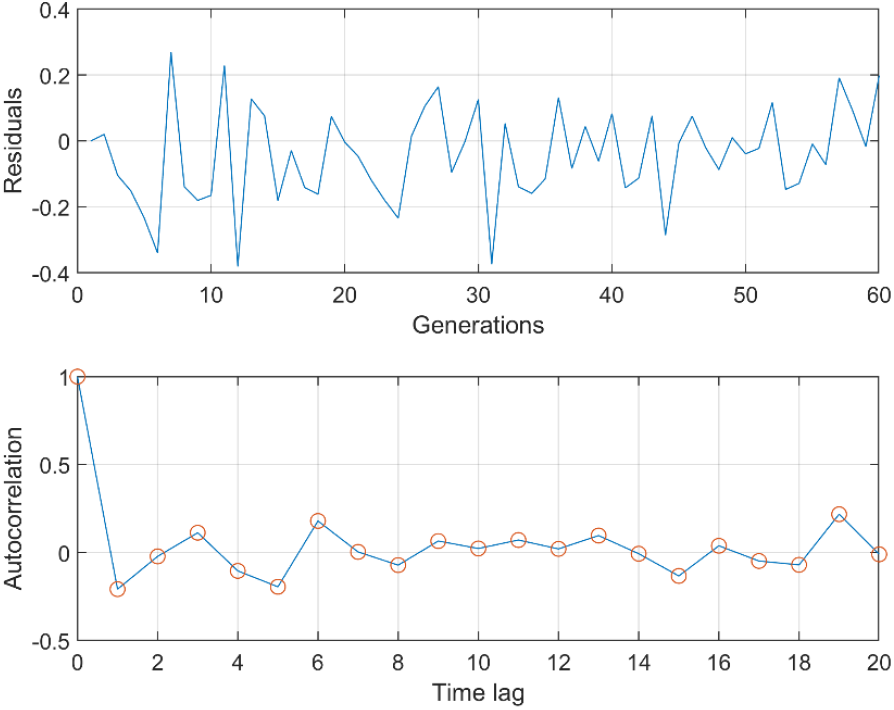
Typical final residuals 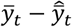 corresponding to the ramp responses in Fig. 5, panels d) to f), and the resulting autocorrelation plot.

#### 3.4.2 Results of errors in *û*_*ref*_

For Case 5 in Table 2, various values of *û*_*ref*_ give prediction errors as shown in Table 3. Note that use of the mean value of the noisy ramp function in Fig. 3, *û*_*ref*_ = 1.25, results in large bias errors in both 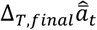 and 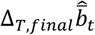. Also note that these bias errors cannot be detected by the proposed validation method, for the simple reason that the modeling and validation data use the same erroneous value of *û*_*ref*_.

**Table 3.**
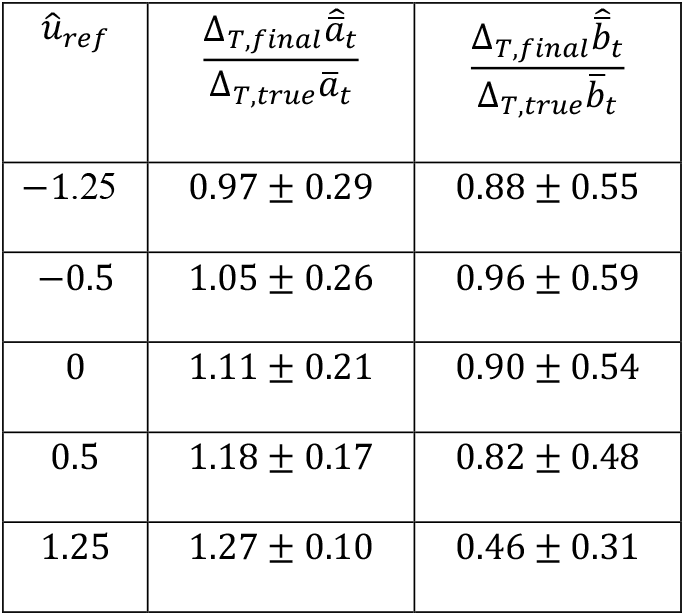
Estimation and prediction results for ramp responses, with guessed values of *u*_*ref*_ and measurement errors in *y*_0,*i*,*t*_, *w*_0,*i*,*t*_ and *u*_0,*t*_ (as in Case 5 in Table 2).

| $\hat{u}_{ref}$ | $\frac{\Delta_{T,final}\hat{\bar{a}}_t}{\Delta_{T,true}\bar{a}_t}$ | $\frac{\Delta_{T,final}\hat{\bar{b}}_t}{\Delta_{T,true}\bar{b}_t}$ |
| --- | --- | --- |
| -1.25 | $0.97 \pm 0.29$ | $0.88 \pm 0.55$ |
| -0.5 | $1.05 \pm 0.26$ | $0.96 \pm 0.59$ |
| 0 | $1.11 \pm 0.21$ | $0.90 \pm 0.54$ |
| 0.5 | $1.18 \pm 0.17$ | $0.82 \pm 0.48$ |
| 1.25 | $1.27 \pm 0.10$ | $0.46 \pm 0.31$ |

#### 3.4.3 Various results

Since an error in *û*_*ref*_ may give a large bias error in 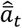 (Table 3), and since 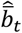 is difficult to predict, it is tempting to set *Ĝ*_*bb*_ = 0, and use the constant estimated slope value, 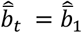. From eq. (6) then follows that 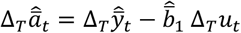, where Δ_*T*_ *u*_*t*_ is independent of *u*_*ref*_. It is also interesting to find prediction results for reduced values of *G*_*aa*_ and *G*_*bb*_. Table 4 summarizes results for various test cases, to be discussed in Section 4. Note that Case 9 gave a very erroneous result for *Ĝ*_*bb*_.

**Table 4.** Prediction results for ramp responses in various test cases with measurement errors in *y*_0,*i*,*t*_, *w*_0,*i*,*t*_ and *u*_0,*t*_. Case 7: *G*_*aa*_ = 0.5 and *G*_*bb*_ = 0.045, and *Ĝ*_*bb*_ = 0. Case 8: *G*_*aa*_ = 0.5 and *G*_*bb*_ = 0.045, and *Ĝ* _*bb*_ = 0 and 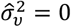. Case 9: *G*_*aa*_ = 0.05 and *G*_*bb*_ = 0.0045, and *Ĝ* _*bb*_ and 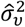 as free variables.

| Test cases | $\hat{G}_{bb}$ | $\hat{b}_1$ | $\frac{\Delta_{T,final}\hat{a}_t}{\Delta_{T,true}\bar{a}_t}$ | $\frac{\Delta_{T,final}\hat{b}_t}{\Delta_{T,true}\bar{b}_t}$ |
| --- | --- | --- | --- | --- |
| Case 7 ($G_{bb} = 0.045$) | 0 | $0.48 \pm 0.02$ | $1.25 \pm 0.21$ | 0 |
| Case 8 ($G_{bb} = 0.045$) | 0 | $0.36 \pm 0.04$ | $2.33 \pm 0.08$ | 0 |
| Case 9 ($G_{bb} = 0.0045$) | $0.022 \pm 0.035$ | $0.49 \pm 0.04$ | $1.10 \pm 0.46$ | $1.72 \pm 2.16$ |

## 4 Summary, discussion and conclusions

Studies of phenotypic responses in wild populations are often based on reaction norm models, where the environmental inputs in many cases are related to climate change. Such input drivers will never be exactly known, and there will also be measurement errors in the recorded responses, and problems related to understanding of ecological mechanisms and prediction of responses are thus error-in-variables problems, as depicted in Fig. 1. Measurement errors in the input signals are especially problematic because they lead to overfitting in parameter estimation, independent of estimation method.

A second important feature of reaction norm response problems is that the environmental inputs must be given an appropriate reference value. This is obvious with an interval scaled input like temperature, but it is just as necessary for a ratio scaled input like salinity. As discussed in Ergon (2022), the proper reference value is the value where the phenotypic variance has a minimum such that geometric mean fitness is maximized, i.e., the environmental value the population is adapted to.

Prediction of evolutionary responses on various ecological drivers must necessarily be accompanied by validation methods (Ljung, 2010; Tredennic et al., 2021), and in order to focus on the essential errors-in-variables and reference environment problems I have here used a reaction norm model in its simplest univariate form, characterized by a mean intercept value and a mean plasticity slope value. I have proposed a validation method where the available input-output data is split into one part for modeling and one part for validation. As shown in simulations in Section 3 and Appendix A, validation of the mean reaction norm trait predictions is difficult due to the short time-series that are normally available from field studies of wild populations. I have also argued that validation of a chosen or estimated reference environment in practice is impossible, when the data comes from the (unintended) anthropogenic global warming experiment, where no independent experimental data with a different reference environment exists.

In simulations for studies of the overfitting and validation problems, I have used a variant of the multivariate breeder’s equation where selection with respect to the unknown individual reaction norm traits is replaced by selection with respect to the individual phenotypic trait, which is assumed to be known from field data (Ergon, 2022). I use a prediction error method (PEM), that is well established in the engineering control field (Ljung, 1998), but it should be noted that overfitting due to errors-in-variables and short data is not a specific problem for PEM, errors in the assumed environmental driver will result in overfitting regardless of the choice of parameter estimation method.

A typical ecological input signal is shown in Fig. 2, where it can be seen that it essentially is white noise on top of a slow trend. Fig. 3 shows the true input signals used in simulations, where the noisy ramp function is similar to typical temperature changes caused by global warming. Fig. 4 illustrates that random errors in ecological drivers lead to overfitting when the squared prediction error is minimized, and that true parameters give better predictions in spite of the fact that they give a larger squared prediction error. This calls for some sort of regularization, but it is unclear how that should be done, if at all possible. Fig. 5 shows typical responses on the noisy ramp input signal in Fig. 3, with added input and output measurement errors, and illustrates that it is easier to predict the mean reaction norm incident than the mean reaction norm slope. Fig. 6 shows that the PEM residuals in the current setting is a white noise sequence, and that PEM thus becomes a maximum likelihood method (Ljung, 2010).

Table 1 shows that parameter estimates and reaction norm predictions with use of the noisy step function in Fig. 3 are quite good also with random output measurement errors (Case 1), but it is then assumed that the reference environment is known. When random input measurement errors are added (Case 2), the results are poorer but still fairly good. Table 2 shows that the noisy ramp input in Fig. 3 with random measurement errors results in large SE values for the mean reaction norm predictions, especially for the mean slope prediction (Case 5). When the number of generations is increased from 60 to 120 (Case 6), the prediction results are clearly improved. The practical problem is thus not only the input measurement errors as such, but the combination with short data. A sufficiently large number of generations is, however, not realistic for many of the studies of wild populations. Table 3 shows that errors in the environmental reference value may give large prediction errors. Mean centering of the ramp function in Fig. 3, will for example give around 30% overestimation of the changes in mean intercept value over 60 generations.

Because input measurement noise makes it difficult to find good and validated predictions of the mean plasticity slope, it is tempting to assume a constant plasticity slope by setting *Ĝ*_*bb*_ = 0. In Case 7 in Table 4 this resulted in 25% overestimation of the change in *ā*_*t*_. The change in *ā*_*t*_ was overestimated also in Cases 2 and 5, but then by only around 10%. A possible explanation of this overestimation is that the measurement error *ũ*_*t*_ is filtered through the non-linear filter given by the prediction model in eq. (4), and that the estimation error increases when *Ĝ*_*bb*_ = 0. There is thus no point in setting *Ĝ*_*bb*_ = 0, and one should definitely not set 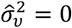, because that results in very large errors in 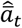 and a large bias in the estimated initial mean slope value 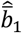 (Case 8 in Table 4). With small values of *G*_*aa*_ and *G*_*bb*_ (Case 9 in Table 4) we find a very large error in *Ĝ*_*bb*_, as further discussed below.

A first main conclusion is that the typically short data from field studies of wild populations make it difficult to predict changes in the mean reaction norm slope value 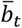, but that the simulation results for changes in the mean intercept value *ā*_*t*_ are somewhat more promising, at least for the fairly large changes in the main examples. The validated ramp response case with input measurement noise (Case 5) gave 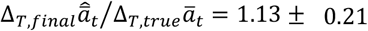 as mean and SE values based on 100 realizations, and four different realizations in Appendix A gave 1.20, 0.99, 1.14 and 0.85. Provided a correct model, the result for a single realization will thus at least give an indication of the change in mean intercept value over all generations. The size of the white input measurement errors used in simulations may be pessimistic, but the simulations assume a perfectly known model structure, which is not realistic, and the input measurements may also have offset errors. It is also promising that the true initial mean slope value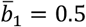, in all normal cases was estimated well, from 0.48 ± 0.02 (Case 5) to 0.50 ± 0.02 (Case 2). The exception is Case 8 in Table 4, where 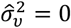 gave 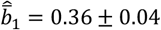.

A second main conclusion is that the proposed validation method is not very reliable for the main cases with *T* = 60 generations. As shown in Appendix A, Examples 1 and 4, there may then occur realizations of ramp responses that give large errors in the mean reaction norm slope predictions, also with seemingly good validation results. This type of problem did not appear when the number of samples was increased to *T* = 120, with *μ*_*U*,*t*_ = 2.5 for *t*> 60.

A third main conclusion is that an assumed or estimated reference environment cannot be validated given data from a specific field study. This is a serious problem since errors in the assumed reference environment may give large prediction errors (Table 3). When studying the effects of the recent global warming it may be tempting to assume that the population was adapted to the mean temperatures in the 1960s, but note that it was around 0.5 °C colder in the mid 1800s, and that evolution often is slow. It should also be noted that estimation of the reference environment by use of the BLUP/PEM optimization method described in Ergon (2023b) gives better results than the gradient method used here, but that this is of limited value without a possibility for validation.

A fourth main conclusion is that small changes in the mean reaction norm traits over all generations result in very unreliable predictions of these changes. Such small changes may be the result of small values of the genetic variances, as in Case 9 in Table 4, where *Ĝ*_*bb*_ was very much larger than the true value. Small changes in the mean reaction norm may also be the result of overlapping generations and long lifetimes, where only a fraction *f*_*t*_ < 1 of the population is affected by the changes in fitness such that the evolution will be slowed down accordingly (Ergon, 2023a). As discussed below, however, small changes in *ā*_*t*_ and 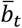 over all generations may also be a result of near optimal adaptive plasticity.

As a short summary of the four main conclusions above, it seems safe to say that disentanglement of plastic and genetic responses on climate change based on realistically short data is a difficult task, and that especially predicted changes in 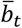 are unreliable, except that the initial mean slope value 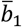 is estimated well. The best option may thus in many cases be to conclude that the mean plasticity slope is constant, 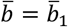, but that does not mean that one should assume that *G*_*bb*_ = 0 in the tuning of the prediction model in eq. (4). As seen in Table 4, Case 7,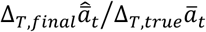 increases to 1.25 ± 0.21 with *Ĝ*_*bb*_ = 0, but this increase will be lower with a smaller variance *G*_*bb*_. There is in any case no point in setting *Ĝ*_*bb*_ = 0 in the search for parameter values, but because of the poor predictions of changes in 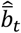 it may be a point in setting 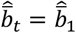.

In the simulations, the plasticity in a stationary stochastic environment is 25% partially adaptive, that is, the plastic phenotypic change is in the adaptive direction, but with only 25% of the optimal magnitude. As shown in Appendix B, an increase to 50% partial adaptivity reduces the changes in *ā*_*t*_ and 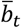 over 60 generations and makes it more difficult to predict these changes. With 100% adaptivity, the initial mean trait values 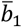 and *ā*_1_ will not evolve when the environment changes, and it will then be impossible to estimate the system parameters. Such an optimal plasticity was an initial assumption in Valdés et al. (2023) (where *G*_*aa*_ and *G*_*bb*_ were far from zero), although the authors eventually found that the plasticity was in fact somewhat larger than optimal (maladaptive plasticity).

The simulations with small values of *G*_*aa*_ and *G*_*bb*_ (Case 9 in Table 4) revealed an identification problem with near 100% adaptivity as discussed above. It is not enough that the system is excited by variations in the environmental input, as shown in Fig. 2, there must also be sufficient changes in *ā*_*t*_ and 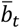 over time. The apparent reason for this can be seen in eq. (1b), 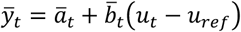, from which follows that when *ā*_*t*_ and 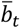 are approximately constant only 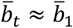 can be estimated from input-output data. The difficulties to estimate *G*_*bb*_ when there are small changes in *ā*_*t*_ and 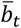 over time raises an interesting theoretical question about persistent excitation in reaction norm identification, but that must be left for further research.

Finally note that I in this article have assumed non-overlapping generations. Field studies of species with overlapping generations and long lifetimes give opportunities to determine individual plasticity, and possibly also to find the reference environment where the phenotypic variance has a minimum. In an example of this approach, Valdés et al. (2023) used 22 years of field observations of the perennial forest herb *Lathyrus vernus* in order to assess phenotypic selection of flowering time as function of spring temperature, as shown in Fig. 2. Although they did not emphasize this result, Fig. 1 in their paper shows that the reference environment where the phenotypic variance has a minimum is the mean temperature in the mid 1800s. This is not surprising in light of the estimated lifetime of 43 years, which means very much overlapping generations and a correspondingly slow evolution.

## A Further examples with 25% adaptive plasticity

Figs. A1 and A2 show four examples of ramp responses with measurement noise in *y*_0,*i*,*t*_, *w*_0,*i*,*t*_ and *u*_0,*t*_ (Case 5 in Table 2), with prediction results given in Table A. Examples 1 and 4 show uncommon but possible extreme responses, which seems to represent good models, but where especially the predictions of 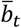 are very poor. This type of responses did not appear when the number of samples was increased to *T* = 120, with *μ*_*U*,*t*_ = 2.5 for *t* > 60.

**Figure A1.**
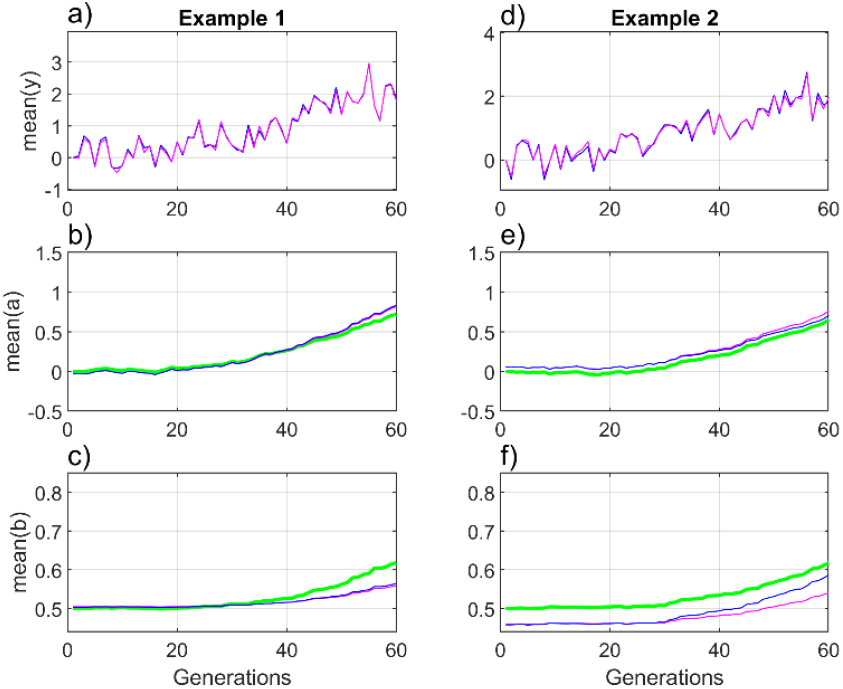
Two examples of ramp responses with 25% partial adaptivity and measurement noise in *y*_0,*i*,*t*_, *w*_0,*i*,*t*_ and *u*_0,*t*_, where Example 1 clearly underestimates *G*_*bb*_ and 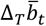 See Fig. 5 caption for details.

**Figure A2.**
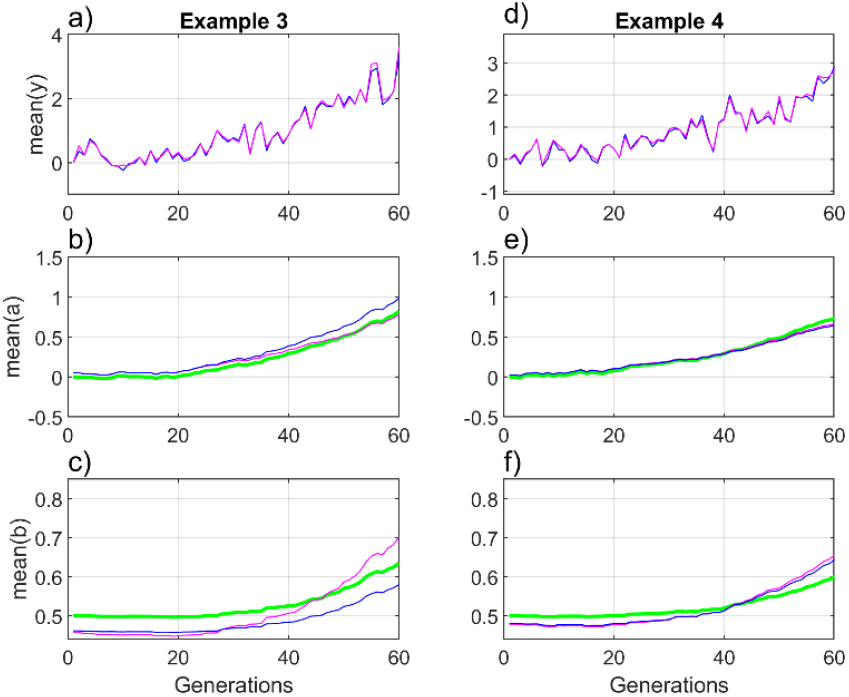
Two additional examples of ramp responses with 25% partial adaptivity and measurement noise in *y*_0,*i*,*t*_, *w*_0,*i*,*t*_ and *u*_0,*t*_, where Example 4 clearly overestimates *G*_*bb*_ and 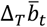. See Fig. 5 captions for details.

**Table A.** Prediction results for the four examples in Figs. A1 and A2.

| Parameters and prediction ratios | True or optimal values | Example 1 | Example 2 | Example 3 | Example 4 |
| --- | --- | --- | --- | --- | --- |
| $\hat{G}_{bb}$ | 0.045 | 0.018 | 0.036 | 0.035 | 0.094 |
| $\hat{\sigma}_v^2$ | 0.5 | 0.39 | 0.49 | 0.39 | 0.57 |
| $\hat{b}_{init}$ | 0.5 | 0.50 | 0.46 | 0.46 | 0.48 |
| $10^4 f_{final}$ | 0 | 42 | 53 | 95 | 52 |
| $\Delta_{T,true} \bar{a}_t / \Delta_{T,true} \bar{y}_t$ | - | 0.39 | 0.36 | 0.25 | 0.26 |
| $\Delta_{T,true} \bar{b}_t / \bar{b}_1$ | - | 0.24 | 0.24 | 0.27 | 0.20 |
| $\Delta_{T,final} \hat{a}_t / \Delta_{T,model} \bar{a}_t$ | 1 | 1.02 | 0.92 | 1.28 | 0.97 |
| $\Delta_{T,final} \hat{b}_t / \Delta_{T,model} \bar{b}_t$ | 1 | 1.12 | 1.59 | 0.49 | 0.92 |
| $\Delta_{T,final} \hat{a}_t / \Delta_{T,true} \bar{a}_t$ | 1 | 1.20 | 0.99 | 1.14 | 0.85 |
| $\Delta_{T,final} \hat{b}_t / \Delta_{T,true} \bar{b}_t$ | 1 | 0.51 | 1.09 | 0.88 | 1.65 |

## B Example with 50% adaptive plasticity

In the system in the simulations in Section 3 and in Appendix A, the environment that maximizes fitness is given by *θ*_*t*_ = 2*u*_*t*_, which implies a linear relationship *μ*_Θ,t_ = 2*μ*_*U*,*t*_ and the variance 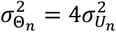. This gives the initial mean plasticity slope 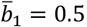, and thus 25% adaptive plasticity. With *μ*_Θ,t_ = *μ*_*U*,*t*_ and 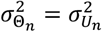, the adaptive plasticity increases to 50%, and as shown in Table B and compared with Case 5, this results in smaller changes of *ā*_*t*_ and 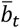 over 60 generations, and larger errors in the predicted changes.

**Table B.** Prediction results for example with 50% adaptive plasticity.

| Parameters and prediction ratios | True or optimal values | Case 5 with 25% adaptive plasticity | Case 5 with 50% adaptive plasticity |
| --- | --- | --- | --- |
| $\hat{G}_{bb}$ | 0.045 | $0.039 \pm 0.030$ | $0.047 \pm 0.043$ |
| $\hat{\sigma}_v^2$ | 0.5 | $0.42 \pm 0.11$ | $0.38 \pm 0.18$ |
| $\hat{b}_1$ | 0.5 | $0.48 \pm 0.02$ | $0.49 \pm 0.03$ |
| $f_{final}$ | 0 | $(73 \pm 37)10^{-4}$ | $(74 \pm 36)10^{-3}$ |
| $\Delta_{T,true}\bar{a}_t/\Delta_{T,true}\bar{y}_t$ | - | $0.34 \pm 0.21$ | $0.16 \pm 0.05$ |
| $\Delta_{T,true}\bar{b}_t/\bar{b}_1$ | - | $0.21 \pm 0.03$ | $0.06 \pm 0.01$ |
| $\Delta_{T,final}\hat{\bar{a}}_t/\Delta_{T,model}\hat{\bar{a}}_t$ | 1 | $1.06 \pm 0.18$ | $1.10 \pm 0.25$ |
| $\Delta_{T,final}\hat{\bar{b}}_t/\Delta_{T,model}\hat{\bar{b}}_t$ | 1 | $351 \pm 1357 *$ | $30 \pm 81 *$ |
| $\Delta_{T,final}\hat{\bar{a}}_t/\Delta_{T,true}\bar{a}_t$ | 1 | $1.13 \pm 0.21$ | $1.19 \pm 0.27$ |
| $\Delta_{T,final}\hat{\bar{b}}_t/\Delta_{T,true}\bar{b}_t$ | 1 | $0.85 \pm 0.52$ | $0.96 \pm 0.84$ |

\*Some values of $\Delta_{T,model}\hat{\bar{b}}_t$ are close to zero.

## C Matlab code

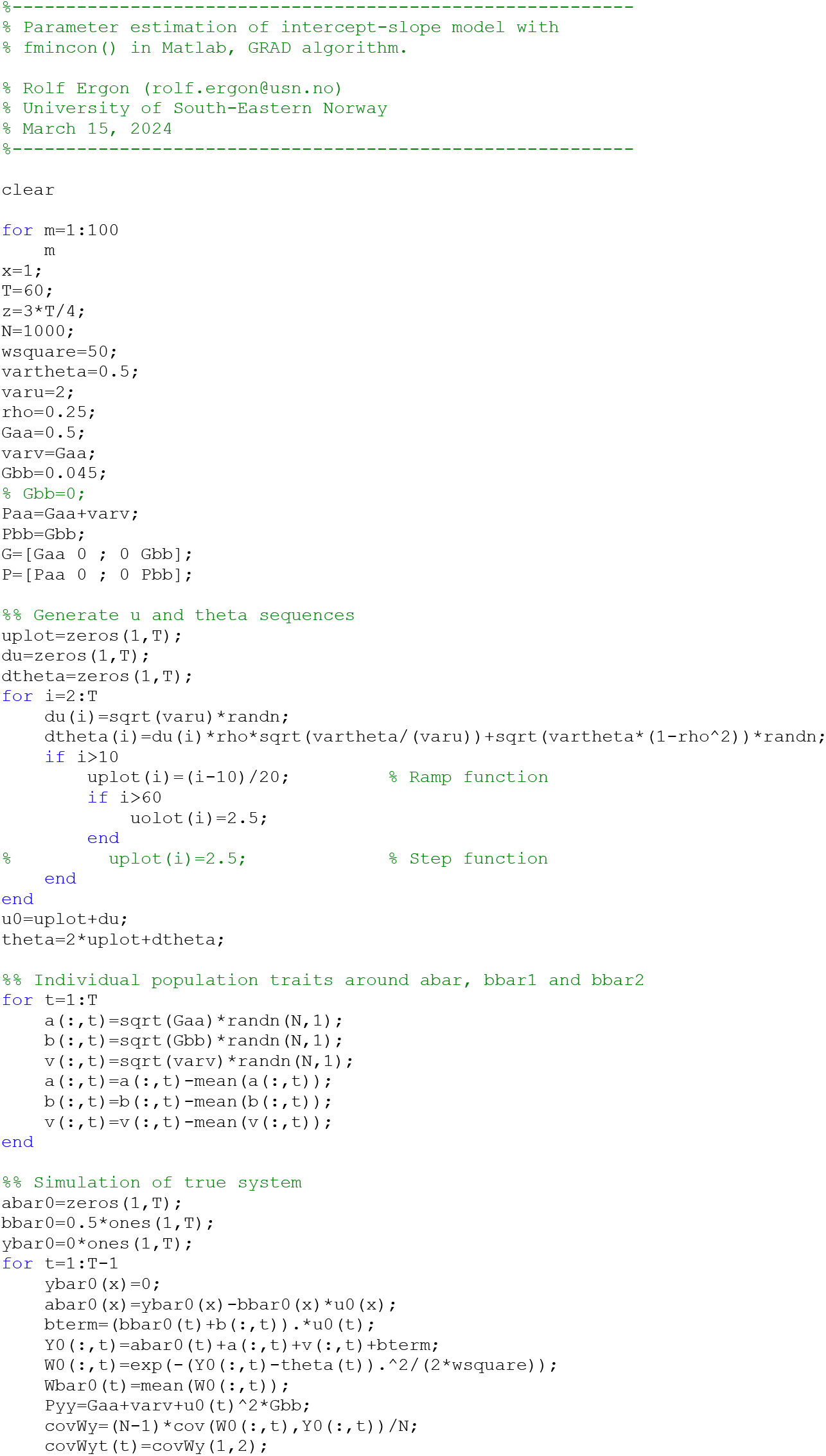
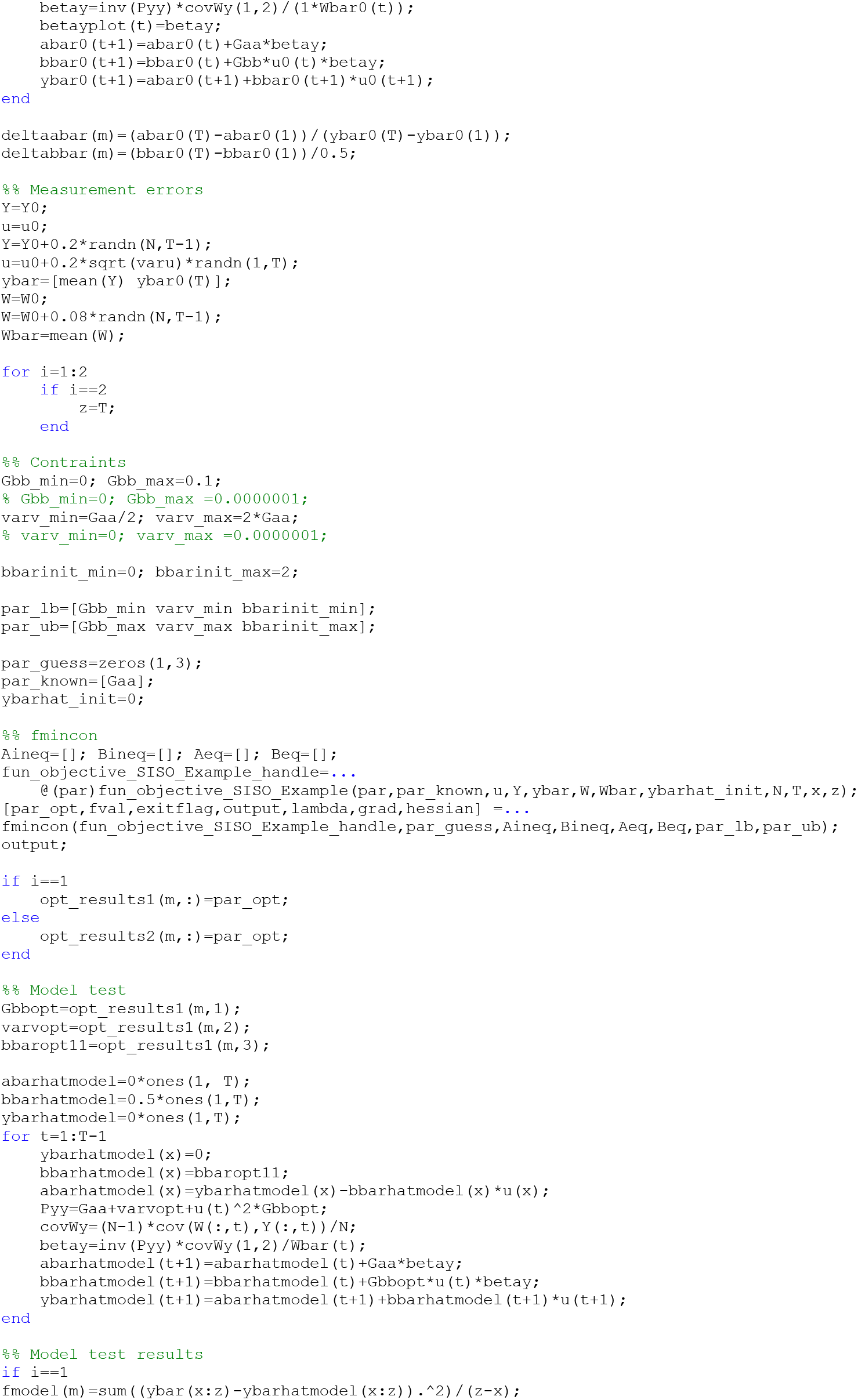
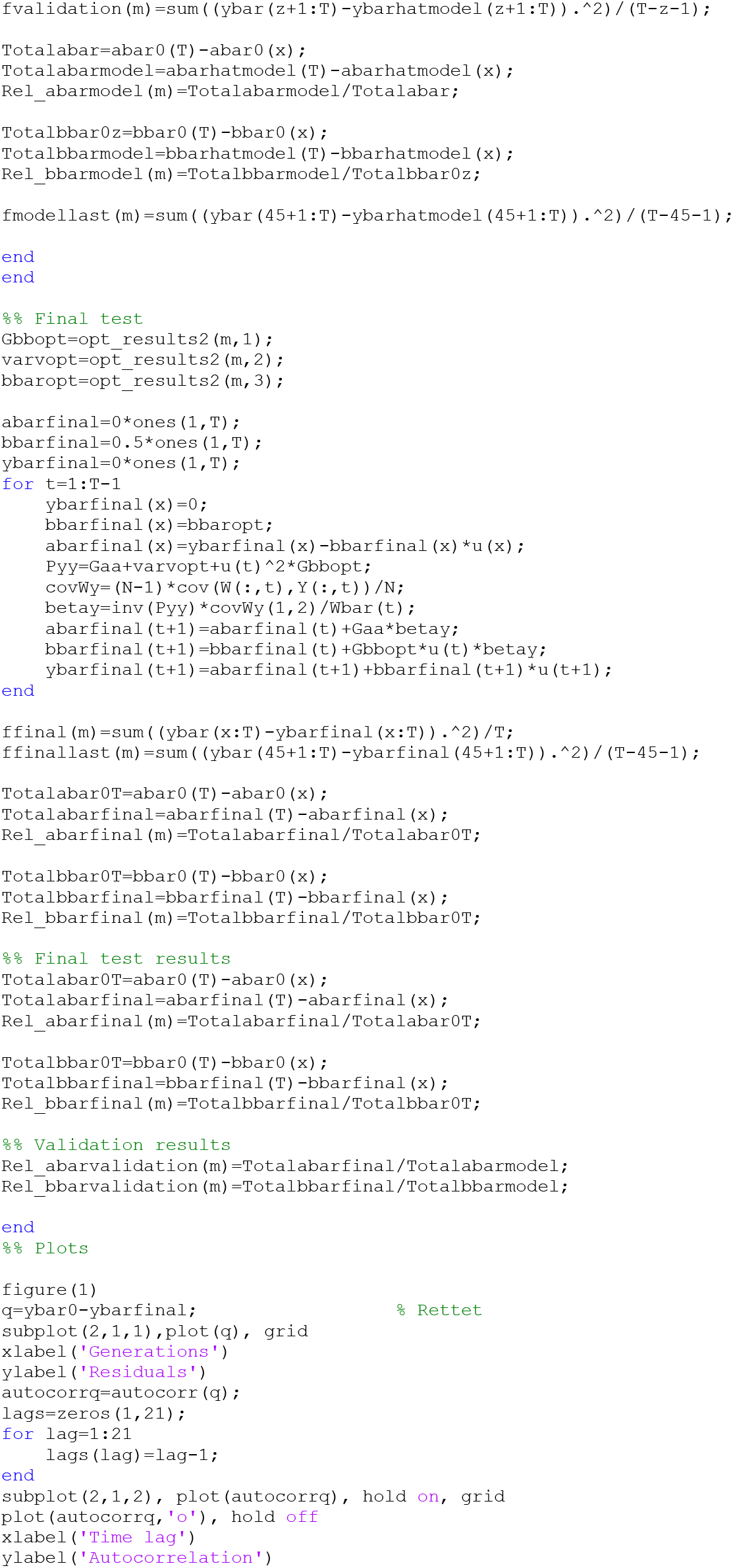
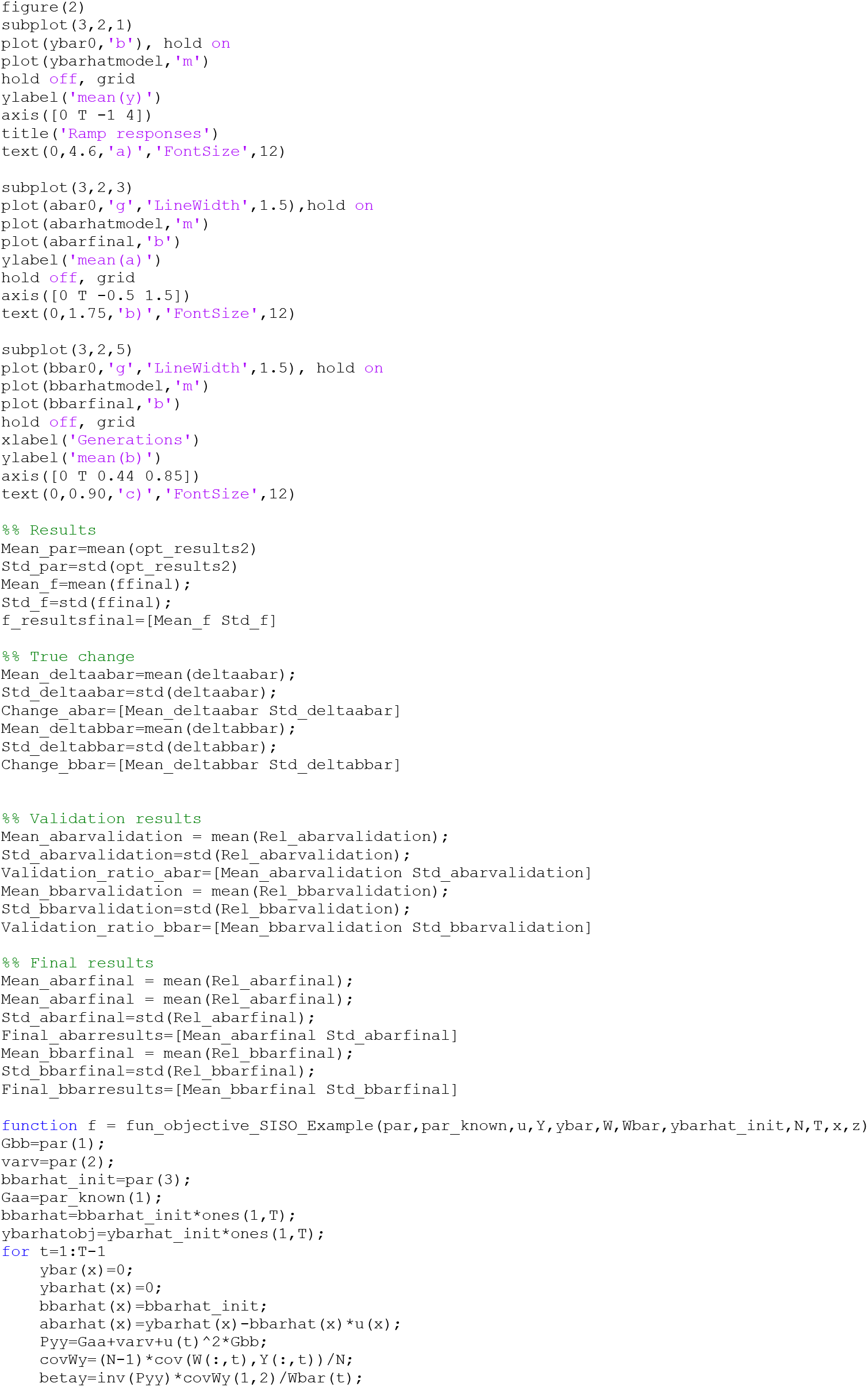
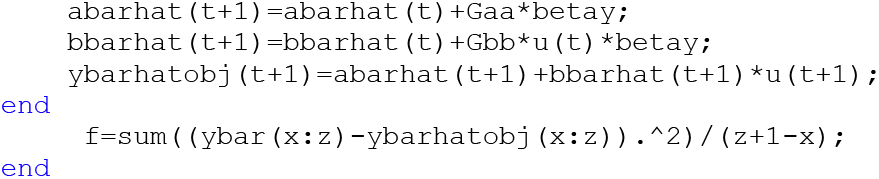

## Notes

### Competing Interest Statement

The authors have declared no competing interest.

